# Lack of negative density-dependence regulation in a dominant oak tree from a neotropical highland forest

**DOI:** 10.1101/329177

**Authors:** Irene Calderón-Sanou, Luis Diego Ríos, Alfredo Cascante-Marín, Gilbert Barrantes, Eric J. Fuchs

## Abstract

Conspecific negative density-dependence(CNDD) is one of the main mechanisms proposed to regulate species coexistence. Tropical highland forests, in contrast to diverse lowland forests, are commonly dominated by a few tree species. Testing the importance of density-dependence effects on seedling establishment of dominant trees may provide insights on the mechanisms regulating population dynamics and forest composition of tropical highlands. We tested the importance of CNDD regulation on seedling survival and recruitment of *Quercus costaricensi*s, a monodominant oak in the Talamanca highland forests of Costa Rica. We used spatial statistics and generalized linear mixed models to test the effects of conspecific density, distance to the nearest adult, density of *Chusquea* bamboo shoots, and herbivory on the annual survival probability of 3538 seedlings between 2014 and 2017. We did not find any effect of CNDD on seedling survival. However, bamboo density and herbivory both significantly decreased seedling survival. All seedlings had signs of herbivory and predator satiation may explain the lack of density dependent regulation in this species. We argue that the lack of intraspecific density regulation at the seedling stage may explain the dominance of *Q. costaricensis* in the highland forests of Costa Rica. Local density of this endemic oak is instead regulated by herbivory and the density of *Chusquea*.

## Introduction

Conspecific negative density-dependence(CNDD) reduces seedling survival and recruitment in many tree species when conspecific density is high. This negative effect likely results from antagonistic intra- and interspecific plant interactions, pathogens and herbivores attracted to high-density patches or through infestation from adults(Janzen 1970; Connell 1971; Freckleton and Lewis 2006; Bagchi et al. 2010; Fricke et al. 2014). Janzen(1970) and Connell(1971) independently proposed that higher offspring mortality of abundant species reduces competition for other species with similar ecological requirements, but susceptible to different enemies, leading to an increase in species diversity. Since then, there has been great interest in testing the importance of this regulating mechanism in both tropical(Clark and Clark 1984; Wright 2002; Peters 2003; Comita et al. 2014) and temperate forests(Hille Ris Lambers et al. 2002; HilleRisLambers and Clark 2003; Yamazaki et al. 2008).

Although most ecologists agree on the ubiquity and importance of CNDD regulation in tropical forests(LaManna et al. 2017), susceptibility to conspecific density may vary across species(Comita et al. 2010; Lin et al. 2012). Rare species are expected to be less affected by CNDD since their populations are small and may not be able to sustain large predator populations(Connell et al. 1984). However, recent findings show that common species are less likely to be affected by CNDD due to a combination of physiological traits and high additive genetic diversity in genes associated with immune responses(Comita et al. 2010; Marden et al. 2017). In tropical plant species, CNDD regulation commonly occurs at the seedling stage and seed-to-seedling transition. Several traits such as a seedling’s shade tolerance or leaf traits(Kobe and Vriesendorp 2011), soil nutrients(LaManna et al. 2016) and diversity and abundance of soil pathogens(Bagchi et al. 2010; Bever et al. 2012), may all influence seedling tolerance to high conspecific densities or to the antagonistic effects of nearby adults(i.e., asymmetric competition).

Tropical forests are commonly perceived as highly diverse habitats, however, tropical montane forests(above 2500 m asl) have fewer species compared to lowland tropical forests and are commonly dominated by a few tree species(Connell and Lowman 1989; Richards 1996; Lieberman et al. 1996). Thereby processes structuring these low diversity tropical forests are not yet fully understood(Hart et al. 1989). In Costa Rica and western Panamá, montane forests are dominated by oaks(Kappelle 2006). *Quercus costaricensis* Liebm.(Fagaceae) is restricted to the Talamanca mountain range, where it represents over 90% of the basal stem area(stems >50 cm dbh) with several thousand juveniles per hectare(Kappelle 2006). Although the dynamics of tropical oak communities have been scarcely studied; temperate oaks have been shown to be susceptible to CNDD and distance to the nearest adult(Wada et al. 2000). In contrast, ectomycorrhizal transfer from adults to close by seedlings seems to have a positive effect on seedling density and survival(Dickie et al. 2002). Therefore, factors regulating seedling recruitment in oaks could be species or population specific, so that conspecific density could either lower survival through competition and herbivory or increase recruitment by sharing symbionts. Considering the pronounced dominance of *Q. costaricensis* in the highlands of Costa Rica and its large density of seedlings, we expected to find weak(if any) CNDD regulation on seedling survival and we expected to find a positive effect of nearby adults on seedling recruitment.

Our main objective in this study was to test for CNDD regulation on survival and recruitment of seedlings of the dominant oak *Q. costaricensis* in the highlands of Costa Rica. Specifically, we tested the effect of conspecific seedling density, distance to nearest adult and herbivory on seedling recruitment and survival. The bamboo *Chusquea* spp.(Poaceae) grows in dense clumps in the understory of montane tropical forests(Widmer 1998) and could be an important factor limiting seedling survival, therefore we also included the effect of bamboo density(*Chusquea* spp.) when studying survival and recruitment of *Q. costaricensis* seedlings.

## Material and Methods

### Study area and species

The study was conducted at Cerro de la Muerte Biological Station(CMBS, 09°33’ N, 83°44’ W; 3100–3350 m a.s.l.), in a Costa Rican upper montane tropical forest(Holdridge 1971). The mean annual precipitation is 2500 mm, with a relatively dry period(<200 mm) between January and April, and the mean air temperatures is 12°C, with large daily fluctuations(from –5 °C to 35 °C)(Fuchs et al. 2010).

The canopy in the study site is dominated by oak trees(*Q. costaricensis*) and the understory by dense clumps of *Chusquea* bamboo(Kappelle et al. 1995; Widmer 1998). *Q. costaricensis* is restricted to upper montane forests(>2200 m elevation) in the Talamanca mountain range in southern Mesoamerica between Costa Rica and Panamá(Hartshorn 1983). Fruiting is synchronous, with a supra-annual masting pattern every two or more years, but flowering occurs every year between December and March(Camacho Calvo and Orozco Vílchez 1998). Acorns fall to the ground between April and October. Seeds germinate after 7 days in controlled conditions, and germination success is 91%(Quirós-Quesada 1990). *Q. costaricensis* hosts both arbuscular- and ecto-mycorrhizal fungi and seedling growth responds positively to these symbionts(Holste et al. 2017). *Quercus* seedlings are highly dependent on cotyledons reserves during early development(Yi et al. 2015) and seed size could have a positive effect in establishment success(Bonfil 1998).

### Seedling censuses

We set four 400-m^2^ plots(20 × 20 m) within CMBS. Plots were placed in old growth forests undisturbed for at least 30 years. Plots were subdivided into 64(2.5 × 2.5 m) quadrats to ease seedling censuses. The density of *Chusquea* stems varied naturally among plots(0 to 14 stems m^−2^). On February 2014, we mapped and tagged all *Q. costaricensis* seedlings (height < 2m) within each plot. We also mapped and measured DBH(> 5 cm) of all adult trees within the plot and in a 10 m wide buffer area around each plot. For each seedling, we measured its height, distance to the nearest conspecific adult and the conspecific seedling density in a 1m radius around each seedling. We also estimated leaf damage by using the qualitative index of herbivory(IH) developed by Núñez-Farfán and Dirzo(1988). For each quadrat, we also counted the number of bamboo stems.

On May/June 2015, 2016 and 2017; we re-censused all plots and determined annual seedling mortality and recruitment per quadrat. A seedling was considered dead when its stem was dry and leafless, rotten or when missing in two consecutive censuses. Dead individuals were removed from survival analyses for the following years. This oak population produced fruits in 2014 and 2016, but less than 15% of adults reproduced in 2016, therefore we considered the 2014 fruiting year as the only masting event during the study period. In 2014, censuses were conducted before acorns had fallen, to avoid including newly germinated seedlings. In 2015, new untagged seedlings were included as the 2015 cohort from the 2014 masting event. These new recruits were easily identified because cotyledons were attached to most seedlings, even after one year. Recruits were mapped and their height and herbivory indexes were also recorded, except in those cases in which seedling leaves were not fully developed(ca. 1% or 15 seedlings). New recruits were included on survival analyses for the following years. Acorns were not produced in 2015 therefore on the 2016 census we only recorded seedling mortality. The few recruits we tallied in 2016(55 in total) were taller than expected for new seedlings, so they were probably uncounted seedlings from the previous census, thus we considered them as 2015 recruits. Only a few adults reproduced in 2016(not a masting year), thus we only recorded five new seedlings in 2017 on all four plots; therefore we only analyzed recruitment success after the 2014 masting.

### Spatial analyses

Both the CNDD regulation and the Janzen-Connell effect depend on seed dispersal and its resulting spatial aggregation. It assumes that seedlings are spatially aggregated. We analyzed the spatial distribution of seedlings in our plots using the L(*t*)=(K(*t*)/π)^0.5^ transformation of the univariate Ripley’s *K*(*t*) function(Ripley 1977), for the 2014 seedlings. We used the R statistical language(Version 3. 2. 2; Core Team 2016) for all spatial analyses(“spatstat” library, Baddeley et al. 2016).

### Survival and recruitment analyses

We modeled the annual probability of seedling survival for three years between 2014 and 2017, as a function of initial height, intraspecific seedling density, distance to nearest conspecific adult tree, herbivory, and *Chusquea* density using a generalized linear mixed-effects model with a *binomial* distribution(Bolker et al. 2009). All variables, as well as the log-transformed seedling height, were standardized and entered in the model as fixed effects. Quadrat and census year were included as random effects. We also included a dummy variable(age) as a fixed factor that identified seedlings from the 2014 census and those recruited in 2015 to determine if newly recruited seedlings had a different survival probability. A second-degree polynomial effect for distance to nearest adult conspecific was also tested. A full model was built including all variables and all second order interactions. To test Janzen-Connell predictions, we analyzed the effect of seedling density and distance to the nearest adult on seedling herbivory(IH). We retain models that included all the variables of interest, significant interactions, and significant polynomial terms (Table S1).

We also analyzed seedling recruitment per quadrat to account for spatial variation in recruitment within a plot. This analysis included only the 2015 data, one year after the masting event of 2014. We used a GLMM with a *quasi-poisson* distribution to estimate the effect of seedling density(number of seedlings per quadrat), conspecific tree density(number of tree trunks in a 2.5 m radius around the center of the quadrat) and *Chusquea* density on seedling recruitment. All explanatory variables were standardized to facilitate comparisons between effects. Plots were included as a random effect to account for spatial autocorrelation of seedlings located within the same plot(Lin et al. 2014). We used the *lme4* package(Bates et al. 2015) in R language(Version 3. 2. 2; R Core Team 2016) for statistical analyses.

## Results

In the 2014 census we mapped and tagged 2031 seedlings, which were spatially aggregated at all scales(Fig. S1). Herbivory was common; over 99% of censused seedlings had some leaf damage and most plants(ca. 71%) had intermediate damage(10-75%), as recorded by the Index of Herbivory(Fig. 1). Contrary to expectations of the Janzen-Connel hypothesis, herbivory decreased at higher seedling densities and increased away from adults(Table S2).

**Fig. 1.**
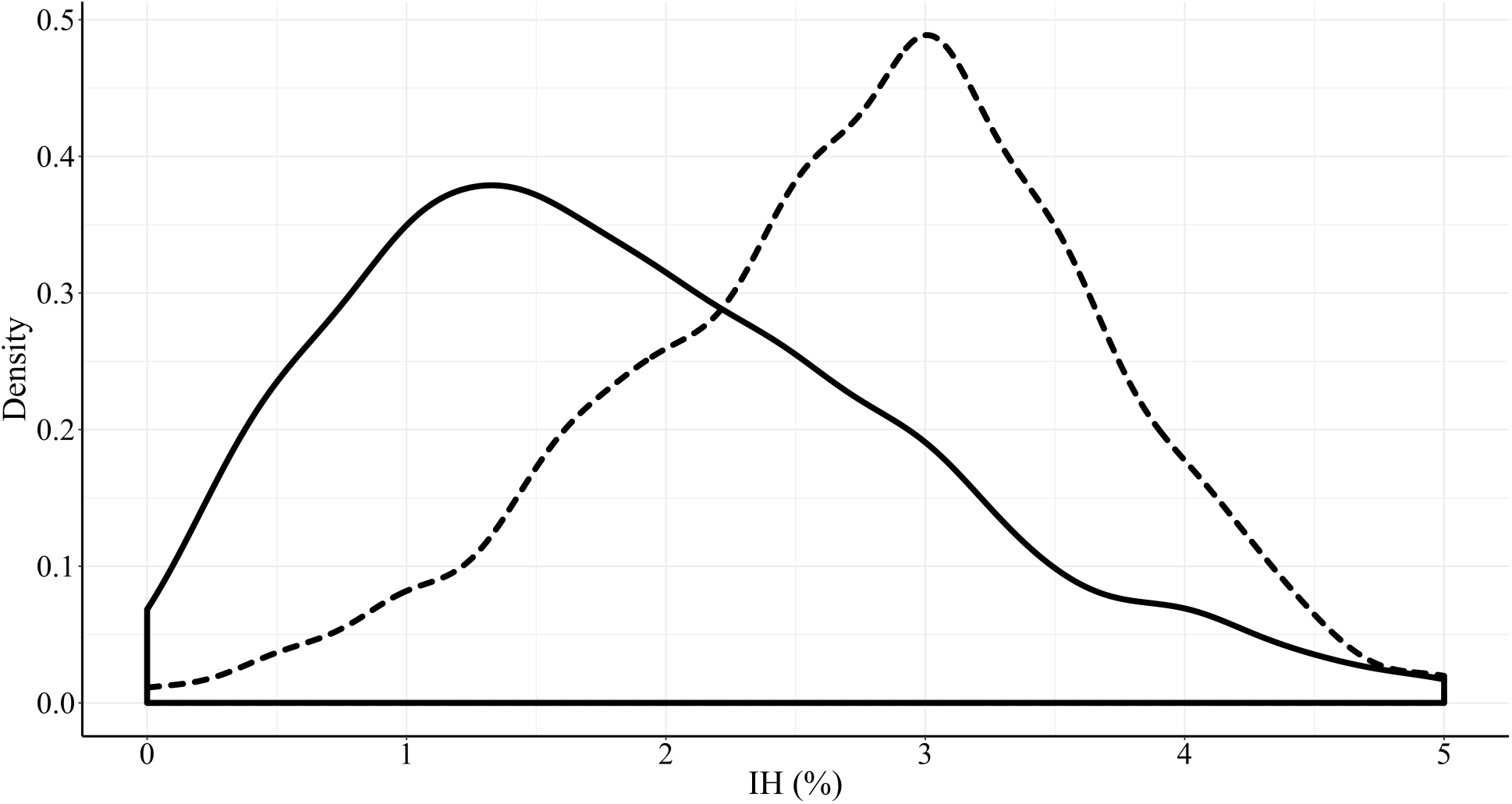
Density distribution of the herbivory index(IH) for seedlings of *Quercus costaricensis*(Fagaceae) in a tropical montane forest in Costa Rica. Dashed line represents seedlings established before the masting event of 2014(N=2031) and solid line, 2015 recruits(N=1578)

Annual seedling survival decreased slightly over the study period, from 73.2% in 2015 to 64.38%(2016) and 63.8%(2017). Seedling height had the strongest positive effect on seedling survival, while bamboo density and herbivory(IH) both reduced survival(Fig. 2). The effect of near adults on seedling survival was best described by a negative polynomial function; seedling survival was highest at intermediate distances from adults(Fig. 2). We found a significant interaction between seedling and *Chusquea* densities: high seedling density significantly increased seedling survival at low bamboo densities(<2.4 stems per m^2^), and decreased survival at higher bamboo densities(Fig. S2). There were no significant interactions between seedling density and herbivory, nor between herbivory and the distance to the nearest adult(Table S1).

**Fig. 2.**
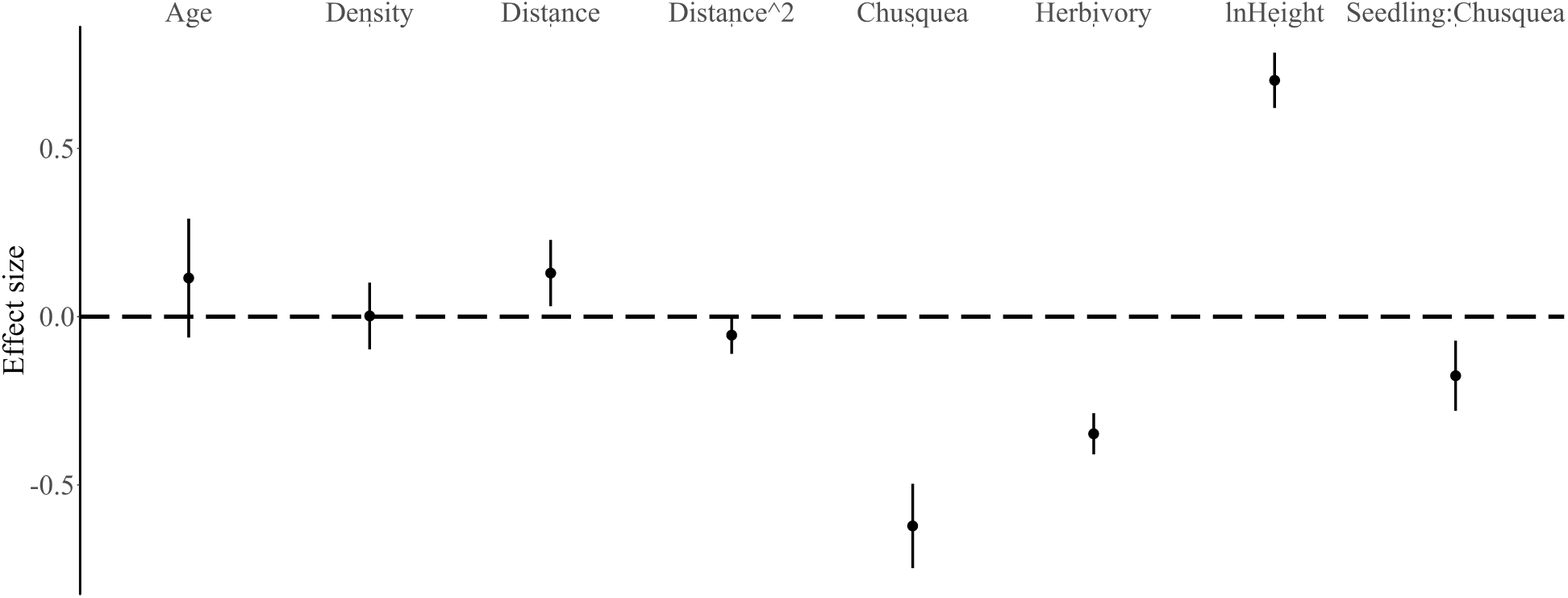
Estimated standardized GLMM effects(±1.96 SE) of factors affecting seedling survival in *Quercus costaricensis* from 2014 to 2017 in a tropical montane forest in Costa Rica. *Age:* describes whether seedlings were censused in 2014 or in 2015; *Density*: intraspecific density within a 1m radius; *Chusquea*: density of bamboo stems; *Distance:* distance to the nearest conspecific adult; *Herbivory:* Herbivory index; *lnHeight:* logarithm of seedling height; *Seedling: Chusquea:* interaction between seedling density and *Chusquea* density

In 2015 we recorded 1578 new recruits. Herbivory was on average lower in new recruits(22.9% leaf damage ± 1.06) compared to seedlings from the 2014 census(43.75% leaf damage ± 0.91; t = 25.798, df = 2828.2, p< 0.0001; Fig.1). Seedling recruitment significantly increased with conspecific seedling density, but the number of new recruits failed to correlate with adult density or with *Chusquea* density(Table S1). Interactions between main factors(seedling, *Chusquea* and tree density) had no effect on seedling recruitment.

## Discussion

Seedlings of the endemic oak species, *Q. costaricensis*, occurred in high density and were spatially aggregated as expected for a masting species with gravity-dispersed seeds(Pulido 2002; Hou et al. 2004). Seedling survival exceeded 63% per year, indicating a relatively high success of seedlings during early recruitment and capacity to survive in dense aggregations. In spite of the densely aggregated distribution of seedlings, we did not detect significant CNDD regulation on seedling survival or recruitment. A similar situation has been reported for other lowland tropical tree species with masting reproduction. For example, several Dipterocarps(Dipterocarpaceae) in Asia(Delissio et al. 2002) or the legume *Dicymbe corymbosa*(Fabaceae) in South America and *Gilbertiodendron dewevrei* in Africa(Hart 1995; Henkel 2003; Henkel et al. 2005); all produce copious amounts of seeds and dense seedling aggregations that escape the negative effects of CNDD regulation; resulting in high adult densities. The lack of negative effects of conspecific density on seedlings establishment of *Q. costaricensis* could contribute to its dominance in Costa Rican montane forests.

Herbivory significantly increased seedling mortality. Loss of leaf area and possible invasion of pathogens through damaged tissue are expected to increase mortality and lower the competitive ability of seedlings(Frost and Rydin 1997). Despite herbivory being ubiquitous in seedlings, survival after three years was high(over 62%) and was unrelated to conspecific density or distance to the nearest adult. These findings disagree with expectations posed by the Janzen-Connell negative density regulation, in contrast, we found a positive effect of density on seedling survival. One possible explanation for the relatively high survivorship of seedlings that we observed, given the high density of oak seedlings(2.4 seedlings m^−2^), could be herbivore satiation(Janzen 1974; Visser et al. 2011; Bagchi et al. 2011). If herbivore satiation occurs, its effect on survival or recruitment is expected to decrease at higher seedling density, when a larger amount of edible tissue is available. Our results support this hypothesis as we recorded higher seedling survival after the masting event(2014-2015, 73.2%) when seedling number increased. We also found a positive effect of density on seedling recruitment. Seeds were more likely to germinate in areas of higher seedling density, where they escape both seed predation and herbivory.

Variability in the relative strength of CNDD regulation between common and rare species has been proposed to have a main effect on determining tree diversity and species composition(Hubbell et al. 2001; Comita et al. 2010). Fricke and Wright(2017) argue that in tropical forests, dominant tree species are expected to have weaker CNDD. The apparent lack of CNDD regulation in *Q. costaricensis* correlates with its high abundance in montane forests and may result from different interacting life history traits(Comita et al. 2010; Zhu et al. 2018). For instance, Kobe and Vriesendorp(2011) found that seedlings from shade tolerant species were less affected by CNDD in lowland tropical forests and suggested that they had better defenses and responses to herbivores. A similar result was found by Zhu et al.(2018) for slow growing tropical species. Abundant species also tend to have low levels of foliar damage that increases individual’s survivorship(Torti et al. 2001). Most seedlings showed low to intermediate levels of herbivory, which does not contribute significantly to CNDD regulation in this Costa Rican oak. High tolerance to leaf damage or long-lived leaves which accumulate minor damage over time, may better explain the observed levels of seedling herbivory.

Ectomycorrizal(ECM) associations increase nutrient acquisition in oak species(Smith et al. 1997), and this may provide an additional advantage to seedling establishment of *Q. costaricensis.* Many dominant tropical trees are associated with ectomycorrhizae and this symbiosis often contributes to their abundance(McGuire et al. 2008). McGuire (2007) found that seedlings from the monodominant tropical tree *D. corymbosa* have a positive input from the ECM network, reducing CNDD effects and increasing seedling survival near parental trees. A balance between asymmetric competition and indirect facilitation by ectomycorrhizal transfer from adults to seedlings increase their survival at intermediate adult densities in *Quercus rubra(*Dickie et al. 2002). Our results show that seedling survival has a negative quadratic relationship with distance to the nearest adult; suggesting that at intermediate distances seedling survival is higher probably due to ECM transfer from adults. At greater distance, ECM colonization may decrease, and competition and CNDD may interact to reduce seedling survival. Given the high density of seedlings near adults, the potential transfer of ECM from adult *Q. costaricensis* may allow seedlings to escape the negative effects of CNDD or mortality through the different mechanisms proposed by Janzen-Connell(Holste et al. 2017).

### The role of interspecific interactions regulating oak population

Interspecific plant interactions play an important role in oak seedling recruitment and survival. The observed aggregated distribution of oak seedlings may be partly determined by localized seed dispersal, but also by differential recruitment influenced by the occurrence of dense patches of *Chusquea* bamboo. The dense thickets formed by bamboo stems strongly reduce oak seedling survival, as they dominate the herbaceous layer limiting the establishment and growth of tree seedlings due to resource competition reducing space available for colonization(Veblen et al. 1979; Royo and Carson 2006; Itô and Hino 2007). The density of bamboo shoots may also significantly increase oak seedling mortality through different mechanisms: resource competition, allelopathy, and physical interference(Veblen 1982; Esteso-Martínez et al. 2006; Royo and Carson 2006). Though *Q. costaricensis* seedlings are shade tolerant(Camacho and Bellefleur 1996), the extreme dense layer of bamboo plants that cover oak seedlings largely reduce the amount of photosynthetic active radiation that reaches the understory, hindering the photosynthetic activity of oak seedlings, and reducing water availability during low precipitation months, since bamboo water intake is very high(Esteso-Martínez et al. 2006). Bamboo could also indirectly interfere with oak seedling development by modifying soil composition; *Chusquea* spp. change the concentration of soil nutrients and litter thickness which may inhibit seedlings establishment or development as observed for *Nothofagus* spp.(Nothofagaceae) in South America(Veblen 1982). *Chusquea* spp. can also hinder the growth of ECM fungi or interfere with their root associations increasing seedling mortality(Wolfe et al. 2008).

In conclusion, the high density of this endemic oak tree in the Costa Rican montane forests is likely the result of predator satiation and adult facilitation through ectomycorrhyzal transference from nearby conspecifics, which results in an apparent absence of CNDD regulation on seedling survival and establishment. Herbivory and interspecific competition with *Chusquea* bamboo appear to be important factors controlling seedling density of this species; but neither of them were density dependent. This lack of CNDD regulation concurs with expectations for dominant tropical trees(Comita et al. 2010). Environmental factors may also influence the strength of CNDD regulation. For example, LaManna et al.(2016) found that the strength of CNDD increases in resource-rich environments. Harsh conditions in montane forests(e.g., acidic soils, low temperatures, high solar irradiance; Kappelle 1996) could impact plant community composition and abundance of natural enemies(McCain and Grytnes 2010), which in turn may alter CNDD regulation(Mangan et al. 2010). However, these factors were beyond the scope of this study.

---

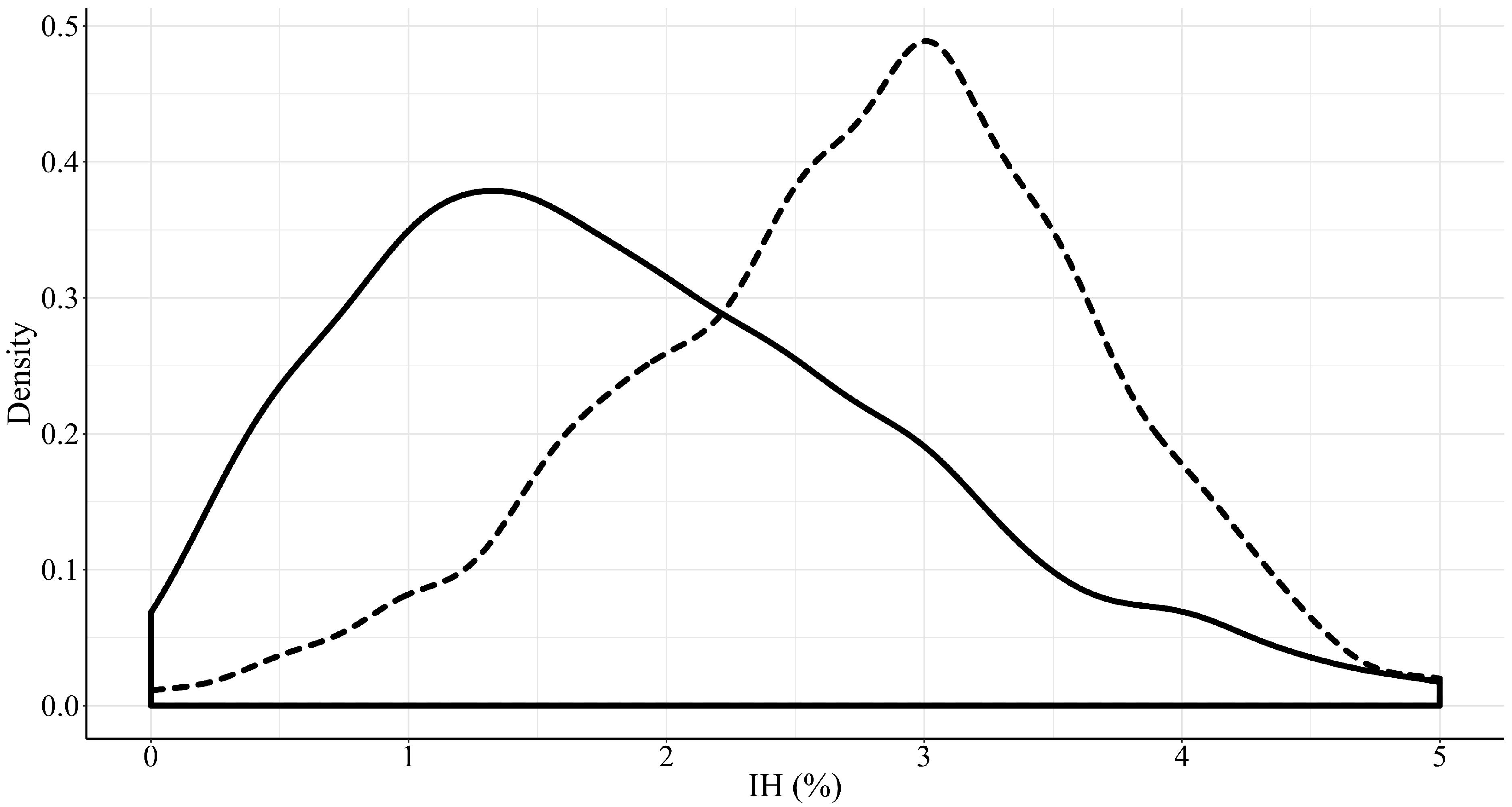

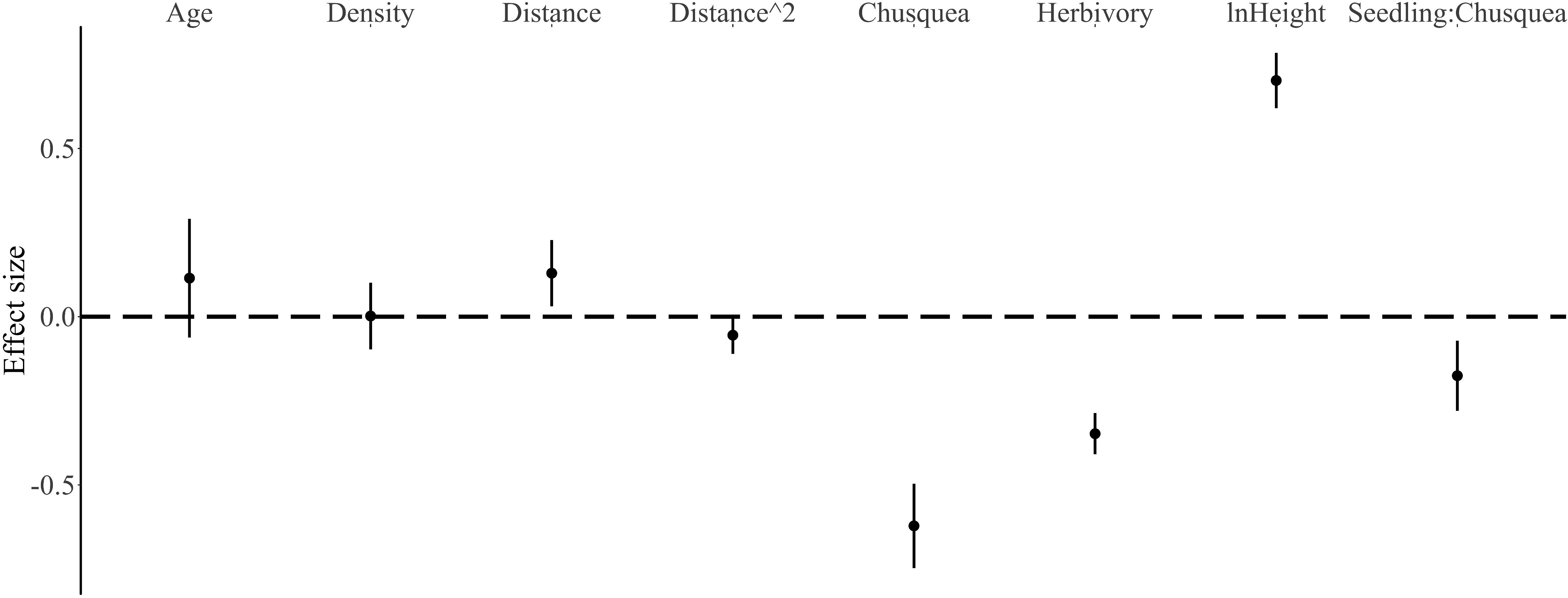

## Acknowledgements

We thank Federico Valverde and Cerro de la Muerte Biological Station for logistic support and Vicerrectoría de Investigación, Universidad de Costa Rica(projects111-B4-087 & 111-B6-A32 to EJF) for financial support. Thanks to all the field assistants that helped make this study possible. This work is part of IC-S undergraduate research project(Licenciatura) at UCR.

